# Dissecting the mechanisms of environment sensitivity of smart probes for quantitative assessment of membrane properties

**DOI:** 10.1101/2022.06.05.494874

**Authors:** Franziska Ragaller, Luca Andronico, Jan Sykora, Waldemar Kulig, Tomasz Rog, Yagmur Balim Urem, Mariana Amaro, Abhinav, Dmytro I. Danylchuk, Andrey Klymchenko, Martin Hof, Ilpo Vattulainen, Erdinc Sezgin

**Author notes:** Deceased (Tomasz Rog).

## Abstract

The plasma membrane serves as a crucial platform for a multitude of cellular processes. Its collective biophysical properties are largely determined by the structural diversity of the different lipid species it accommodates. Therefore, a detailed investigation of biophysical properties of the plasma membrane is of utmost importance for a comprehensive understanding of biological processes occurring therein. During the past two decades several environment-sensitive probes have been popular tools to investigate membrane properties. Although these probes are assumed to report on membrane order in similar ways, their individual mechanisms remain to be elucidated. In this study, using model membrane systems, we studied the probes Pro12A, NR12S and NR12A in depth and examined their sensitivity to parameters with potential biological implications, such as the degree of lipid saturation, double bond position and configuration, phospholipid headgroup and cholesterol content. Applying spectral imaging together with atomistic molecular dynamics simulations and time-dependent fluorescent shift analyses, we unravelled individual sensitivities of these probes to different biophysical properties, their distinct localizations and specific relaxation processes in membranes. Overall, Pro12A, NR12S and NR12A serve together as a toolbox with a wide range of applications allowing to select the most appropriate probe for each specific research question.

## Introduction

Besides functioning as a barrier, the plasma membrane is involved in a multitude of cellular processes such as cell division, endo- and exocytosis, intracellular membrane trafficking and cell signalling [1]. To carry out all its functions, the plasma membrane is highly heterogeneous, harbouring numerous membrane-associated proteins, lipids, proteoglycans, glycolipids, and glycoproteins [2]. The lipid bilayer itself also comprises hundreds of structurally diverse lipid species [3] heterogeneously distributed across and within each bilayer leaflet [4].

The structural diversity of the different lipid species defines the collective biophysical properties of the membrane such as membrane fluidity, tension, curvature, phase transition, and charge distribution [4,5]. Cellular processes can evoke changes in local membrane composition and consequently the biophysical properties of the membrane are constantly altered, allowing functional adaptation of the membrane [2,6]. To gain a more detailed in-depth understanding of the processes occurring at the plasma membrane and their regulation, studying its biophysical properties is of utmost importance [5]. This highlights the need for developing new tools of investigation.

During the past two decades several fluorescent probes have been developed to study different biophysical properties of biomembranes [7]. Environment-sensitive probes (smart probes) have become particularly popular as they can sense membrane properties such as polarity, viscosity, hydration and tension, among others. These probes change their emission wavelengths, intensity or fluorescence lifetime depending on their immediate environment [8]. As a classical example for a solvatochromic probe, Laurdan can distinguish between liquid ordered (Lo; saturated lipid phase) and liquid disordered (Ld; unsaturated lipid phase) membranes by a red-shift in the emission wavelength [9,10]. Another example for a solvatochromic probe is di-4-ANEPPDHQ, which exhibits a strong spectral shift depending on polarity of the immediate environment [11]. Being both polarity-sensitive probes, they were often assumed to report on the same properties of membranes; however, recent work revealed that Laurdan and di-4-ANEPPDHQ report on different properties of membranes [12]. This discrepancy in the mechanism of spectral shift highlights a new potential opportunity: can we use different polarity-sensitive probes to measure quantitatively different properties of membranes, such as lipid saturation index, configuration and position of lipid unsaturation, lipid headgroup, and cholesterol content?

Laurdan and di-4-ANEPPDHQ have certain disadvantages, hampering their widespread use in cell biology [12,13]. The two main drawbacks of Laurdan are poor photostability and high internalization [13], whereas di-4-ANEPPDHQ exhibits complex photophysics (i.e. sensitivity to multiple biophysical properties) [12]. Moreover, both probes require relatively high concentration for cell membrane staining. Ideally, environment-sensitive solvatochromic probes for plasma membrane should *i)* be fluorogenic and bright (high signal-to-noise ratio); *ii)* exhibit a strong fluorescence dependence on the environmental properties; *iii)* exclusively localize in the plasma membrane with low internalization and toxicity; and *iv)* be suitable for live-cell imaging to study membranes at physiological conditions. More recently, new environment-sensitive probes were designed to fulfil these criteria and to overcome previous limitations, such as Pro12A (a derivative of Laurdan) [14], NR12S and NR12A (derivatives of Nile Red) [15,16]. Compared to Laurdan and Nile Red, Pro12A, NR12S and NR12A bear an anchor group composed of a charged (anionic sulfonate or zwitterionic) head group with a lipophilic alkyl chain and can thereby partition almost exclusively into the outer leaflet of the plasma membrane [14–16]. Compared to Laurdan, Pro12A is brighter, shows a larger spectral shift between ordered and disordered phases, and minimized internalization [14]. NR12S exhibits a strong spectral shift towards the red between ordered and disordered phases, and hardly shows any internalization compared to Nile Red [16]. NR12A, which has an inverted Nile Red moiety compared to NR12S, additionally exhibits improved brightness and photostability compared to NR12S [15].

These probes are poised to open new avenues for the investigation of cellular membranes; however, their mechanism of spectral shift should be thoroughly investigated to be able to fully exploit them. The spectral shift of fluorescent probes due to an environmental change is usually quantified by calculation of intensity ratios or generalized polarization (GP) parameter. High ratios and GP-values are correlated with higher lipid order, increased lipid packing, and a decrease in polarity in the membrane [17,18]. However, intensity ratios and GP-values cannot directly report on the underlying biophysical property causing the spectral shift of the fluorescent probe. Therefore, it remains to be elucidated which biophysical property (or properties) the fluorescent probes are mainly sensitive to. To evaluate this, we characterized the three recently developed smart-probes Pro12A, NR12S and NR12A in depth. We examined their sensitivity to parameters with potential biological implications, such as the degree of lipid saturation, double bond position and configuration (*cis* vs. *trans*), phospholipid headgroup, and cholesterol content. Thereby, we screened for one parameter at a time in spectral imaging and subsequent GP-analysis, by using model membrane systems with controllable complexity. To understand the impact of the smart probes’ orientation and localization in the membrane on the observed discrepancies in sensitivity to different properties, we carried out *in silico* atomistic molecular dynamics (MD) simulations. These biomolecular simulations provided significant added value in understanding experimental data [19–22]. Furthermore, we performed time-dependent fluorescence shift (TDFS) experiments to investigate the mechanism and dynamics of the smart probes’ spectral shift. Overall, the results reveal that Pro12A performs particularly well at sensing cholesterol content, while NR12S is superior in differentiating the degree of saturation and the phospholipid headgroup, and NR12A is best suited to differentiate between varying positions and configurations of the double bond in unsaturated lipids. These smart probes adopt varying orientations and locations within the membrane, which is in line with observed sensitivities. Further, TDFS analysis indicates that the smart probes sense changes in fluidity.

In summary, these environment-sensitive probes provide a useful tool to investigate different aspects of the plasma membrane in cellular processes. Our in-depth analysis of the smart probes offers great applications in biology by providing insights on how to choose the right probe best suited for the investigation of the membrane property of interest or combine them to obtain a more holistic picture of biophysics of biomembranes.

## Materials and Methods

### Materials

We used the following lipids and environment-sensitive probes: 1,2-diarachidonoyl-sn-glycero-3-phosphocholine (DAPC, 20:4/20:4 PC), 1,2-dipetroselenoyl-sn-glycero-3-phosphocholine (Δ6cis DOPC, 18:1/18:1 PC), 1,2-dioleoyl-sn-glycero-3-phosphocholine (Δ9cis DOPC, 18:1/18:1 PC), 1,2-dielaidoyl-sn-glycero-3-phosphocholine (Δ9trans DOPC, 18:1/18:1 PC), 1-palmitoyl-2-oleoyl-glycero-3-phosphocholine (POPC, 16:0-18:1 PC), 1,2-dipalmitoyl-sn-glycero-3-phosphocholine (DPPC, 16:0/16:0 PC), 1-palmitoyl-2-oleoyl-sn-glycero-3-phospho-L-serine (POPS, 16:0/18:1 PS), 1-palmitoyl-2-oleoyl-sn-glycero-3-phosphoethanolamine (POPE, 16:0/18:1 PE), cholesterol (Avanti Polar Lipids), Pro12A [14], NR12S [16] and NR12A [15].

NaCl, HEPES, CaCl_2_, ethylenediaminetetraacetic acid (EDTA) and dithiothreitol (DTT) were obtained from Sigma-Aldrich (St. Louis, MO, USA). Paraformaldehyde (PFA) was obtained from ThermoScientific, and PBS and high-glucose DMEM from ThermoFisherScientific. Organic solvents of spectroscopic grade were supplied by Merck (Darmstadt, Germany).

### Large Unilamellar Vesicle Preparation

LUVs of the following compositions were prepared: Δ9*cis* DOPC, POPC, POPC:Chol 90:10, POPC:Chol 50:50 and DPPC:Chol 50:50. Chloroform solutions of the lipids were combined in the appropriate amounts. The solutions were then mixed with a methanol solution of the fluorescent probe to a final molar ratio of lipids to probe of 100:1. The organic solvents were evaporated under a stream of nitrogen. For thorough removal of the solvent, the lipid films were left under vacuum for at least 1.5 h. Buffer (10 mM HEPES, 150 mM NaCl, 0.2 mM EDTA, pH 7.1) was then added to the dried lipid film (lipid concentration of 1 mM) and each sample was vortexed for one minute and 6 freeze-thaw cycles were performed using liquid nitrogen and a water bath at 60 °C. Samples were prepared via extrusion through a polycarbonate membrane with a nominal pore diameter of 100 nm (Avestin, Ottawa, Canada) to yield the final LUV suspensions. POPC:Chol 50:50 and DPPC:Chol 50:50 extrusions were performed at 60 °C. For measurements the vesicle suspension was diluted to an overall lipid concentration of 0.5 mM.

### Giant Unilamellar Vesicle (GUV) Preparation

GUVs were prepared according to a previously described protocol [23]. GUVs of the following lipid compositions were prepared: DAPC, Δ6cis DOPC, Δ9cis DOPC, Δ9trans DOPC, POPC, POPC:Chol 90:10, POPC:Chol 80:20, POPC:Chol 50:50, DPPC:Chol 50:50, POPC:POPS 90:10, POPC:POPE 90:10 and SM:DOPC:Chol 2:2:1 (phase-separated GUVs). In short, GUVs were generated by electroformation using custom-built GUV Teflon chambers with two platinum electrodes [24]. A volume of 6 µl of lipid dissolved in chloroform (1 mg/ml total lipid concentration) was homogeneously distributed on the electrodes, dried under nitrogen stream and placed in 300 nM (370 µl) sucrose solution. Electroformation was performed at 2 V and 10 Hz for 1 h, followed by 2 V and 2 Hz for 30 min. GUV preparation of DPPC:Chol or SM:DOPC:Chol was carried out above the specific lipid transition temperature at 70 °C, whereas the other GUVs were generated at room temperature. To confirm successful integration of POPS in POPC:POPS 90:10 GUVs after electroformation, 1 µl of Annexin V-Alexa Fluor 647 (Thermo Fisher Scientific) was added to 100 µl GUVs.

### Cell Maintenance and Giant Plasma Membrane Vesicle Preparation

U2OS cells were cultured in DMEM (high glucose, without pyruvate) with 10 % FBS at 5 % CO_2_ and 37 °C. They were seeded in 6-well plates to reach 50-60 % confluency on the day of GPMV production. GPMVs were prepared according to a previously described protocol [23]. In short, after washing the seeded cells (50-60 % confluency) twice with GPMV buffer (150 mM NaCl, 10 mM HEPES, 2 mM CaCl_2_, pH 7.4), the cells were incubated in 1 ml GPMV buffer with 25 mM PFA and 50 mM DTT (final concentrations) at 37 °C for 2 h before collection of the GPMVs in the supernatant.

### Membrane labelling and Confocal Spectral Imaging

GUVs and GPMVs were stained at a final concentration of 100 nM of NR12S or NR12A, and 300 nM Pro12A. The GUVs and GPMVs were imaged in previously blocked (3 mg/ml BSA in PBS) µ-Slides (18 well glass bottom, ibidi) at room temperature.

For Confocal Spectral Imaging of GUVs and GPMVs, a Zeiss LSM 780 confocal microscope with a 32-channel array of gallium arsenide phosphide (GaAsP) detectors was utilized. Pro12A was excited at 405 nm and NR12S and NR12A at 488 nm. The emitted fluorescence was collected simultaneously in ∼9 nm wavelength intervals between 423 and 601 nm for Pro12A (20 channels), or between 503 and 700 nm for NR12S and NR12A (22 channels). To obtain the spectra from the intensity values of all detection channels, a region of interest of similar size was selected for the liquid ordered and disordered phases of a GUV or GPMV as well as for the background and a Z-profile was generated using ImageJ. The intensity values of liquid-ordered and disordered phases were corrected for the background and then normalized. This was performed for three phase-separated GUVs and GPMVs stained for each probe.

### GP Analysis

The Generalized Polarization (GP) value is calculated using Equation 1:

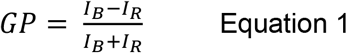

where *I*_*B*_ and *I*_*R*_ are the fluorescence signal intensities at the blue- or red-shifted emission wavelengths, respectively, for liquid ordered, λ_Lo_, and liquid disordered phase, λ_Ld_. According to Equation 1 the GP can adopt values ranging between +1 and -1. We defined λ_Lo_ and λ_Ld_ as 423 and 494 nm for Pro12A, 557 and 664 nm for NR12S, and 583 and 673 nm for NR12A. At these wavelengths the differences in intensity of ordered and disordered phases were high and led to the largest GP range, while not using channels with too few fluorescence signals. Different wavelengths for λ_Lo_ and λ_Ld_ can be chosen, which might increase the GP contrast. The GP value is a relative value depending on λ_Lo_ and λ_Ld_ as well as on the environment-sensitive probe.

### Data analysis script

The GP of vesicles was measured using a homemade Python script built from common packages (i.e., *NumPy, scikit-image, scikit-learn*, and *matplotlib*). Briefly, automatic thresholding (via Otsu algorithm) was performed on each image to generate a mask. On the latter, a density-based clustering algorithm (DBSCAN) allowed to isolate single vesicles from clusters according to the respective object-detected eccentricity value (i.e., objects with eccentricity ≥ 0.5 were discarded). Then, pixel-wise GP values of individual vesicles were calculated according to Equation 1 to estimate the respective GP median and standard deviation. For each dye, a different pair of wavelengths were selected to measure the fluorescence intensity from the ordered (Lo) and disordered (Ld) phase, namely 423 nm (Lo) & 494 nm (Ld) for Pro12A, 557 & 664 nm for NR12S and 583 & 673 nm for NR12A. The GP profile of individual vesicles (see Figure 1H-J) was obtained by calculating the GP value of an area 3×1 pixels^2^ (integrating element) sliding along a line centred onto the membrane.

**Figure 1:**
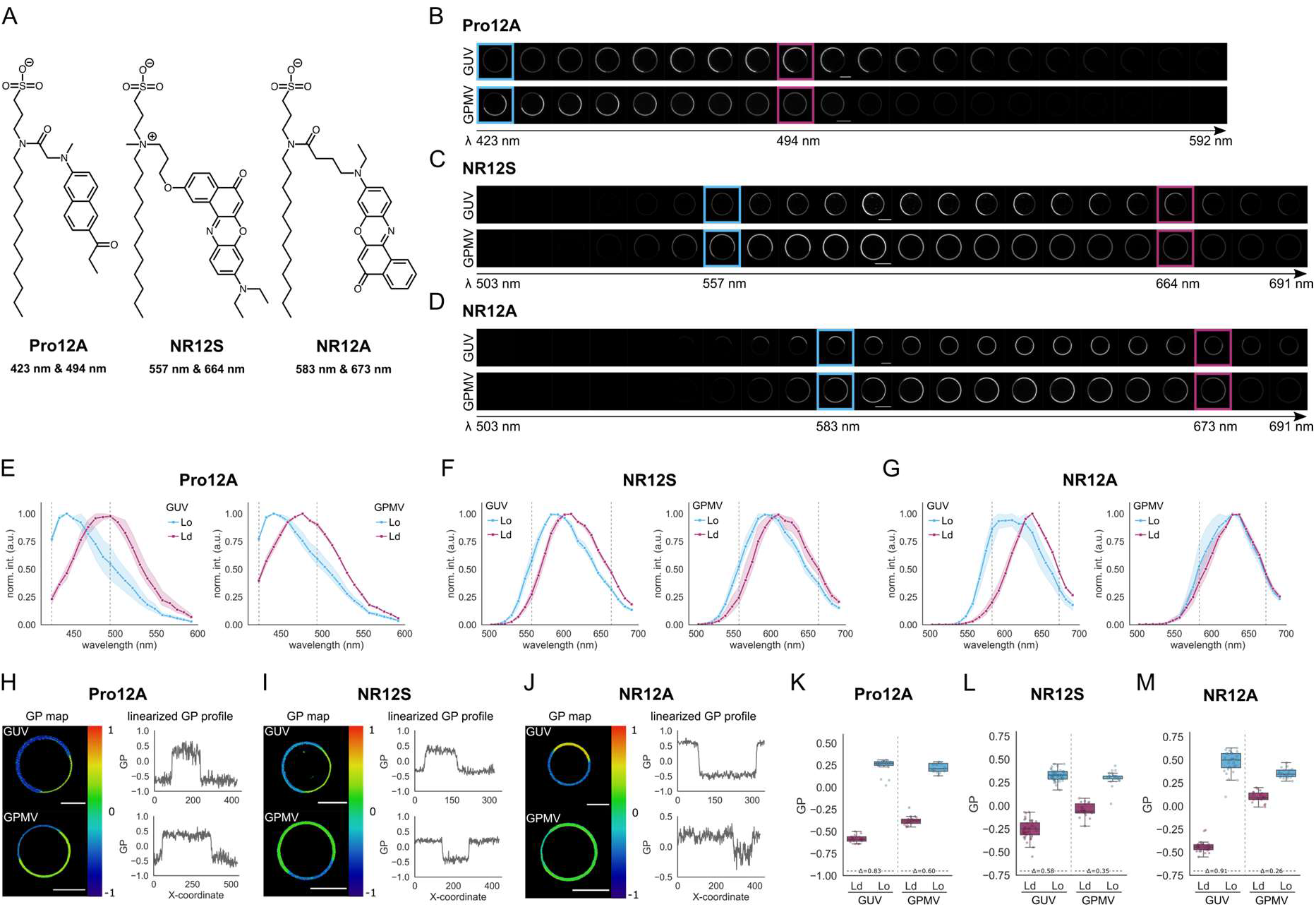
Spectral shift of Pro12A, NR12S and NR12A in phase-separated GUVs and GPMVs. Spectral imaging overview of Pro12A, NR12S and NR12A. A| chemical structure of the probes Pro12A, NR12S and NR12A. B-D| Array of spectral images obtained from the spectral detector (∼9 nm intervals) of GUVs and GPMVs stained with Pro12A (B), NR12S (C) or NR12A (D). Coloured boxes indicate the channels that were used for GP calculation. E-G| Normalized intensity spectra of Pro12A (E), NR12S (F) or NR12A (G) in GUVs (left) or GPMVs (right). Ordered phase is shown in blue and disordered phase in magenta. Squares mark the average and the surrounding band the standard deviation calculated from three vesicles. Dashed lines correspond to the wavelength-channels used for GP calculation. H-J| Example images of GP-colour coded GUV and GPMV (left) and linearized GP profiles along the vesicles (right) stained with Pro12A (H), NR12S (I), NR12S (J). K-M| Calculated GP values of ordered and disordered phases in GUVs and GPMVs stained with Pro12A (K), NR12S (L) and NR12A (M). Value between dashed line corresponds to the ΔGP values calculated from the mean. One representative replicate of three is shown for GUVs and GPMVs.

### Statistics

Statistical analysis was performed for the median GP of the GUVs of all 3 replicates taken together. Using the Python library *SciPy* with its module *scipy*.*stats*, a Kruskal Wallis test was performed to test for significance (α=0.05) within a group (saturation, configuration & position of the double bond, headgroup and cholesterol content). Using the package *scikit-posthocs*, post-hoc Mann-Whitney U tests (with Bonferroni correction) were performed to test for significance (α=0.05) between individual lipid compositions within one group.

### LUV spectroscopy and time dependent fluorescent shift measurements

Steady-state excitation and emission spectra were acquired using a Fluorolog-3 spectrofluorometer (model FL3-11; Jobin Yvon Inc., Edison, NJ) and FS-5 spectrofluorometer (Edinburgh Instruments, U.K.) equipped with a xenon arc lamp. The temperature in the cuvette holders was maintained using a water-circulating bath. The steady-state spectra were recorded in steps of 1 nm (bandwidths of 1.2 nm were chosen for both the excitation and emission monochromators) in triplicates and averaged.

Fluorescence decays were recorded on a 5000 U single-photon counting setup (IBH, Glasgow, U.K.) using a laser excitation source (peak wavelengths of 375 nm (LDH-DC-375, Picoquant, Germany) and 532 nm (PicoTa, Toptica Photonics AG, Germany), for Pro12A and NR-probes, respectively, at a 5 MHz repetition rate) and a cooled R3809U-50 microchannel plate photomultiplier (Hamamatsu, Japan). A 399 nm or 550 nm cut-off filter was used to eliminate scattered light. The signal was kept below 1 % of the repetition rate of the light source. Data were collected until the peak value reached 5000 counts. Measurements were performed at 23°C and 37 °C.

### Estimation of fluorescence spectrum at time zero, ν *(0)*

Deep temperature emission and excitation scans were performed for Pro12A and NR12A in Ethanol:Methanol mixture (1:1) forming transparent glass. The spectra were recorded using Fluorolog-3 spectrofluorometer (model FL3-11; Jobin Yvon Inc., Edison, NJ) equipped with dewar liquid nitrogen accessory at approximately 90 K. The fluorescence spectra at “time zero” were determined as the emission maxima of the deep temperature scans corrected for the varying maxima of the excitation spectra between the deep temperature scans and those recorded in lipid systems.

For the samples with NR12S dye the deep temperature scans did not yield acceptable results and thus a different approach was taken where the simplified procedure for the “time zero” estimation, ν*(0)*, was used [25]:

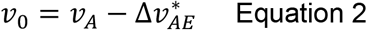

where ν_*A*_ is the maximum of the absorption spectrum in the system of interest, *Δ*ν_*AE*_*\** is the value gained by extrapolation of the dependence of the dye’s stokes shift, *Δ*ν_*AE*_, on the polarity function *Δf* when *Δf* → 0. For this the Lippert-Mataga plot [26–28] was used, where the polarity functions, *Δf*, is defined as:

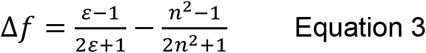

Where *ε*_*r*_ is the dielectric constant and *n* the refractive index of the solvents.

### TDFS Analysis

Fluorescence emission decays were recorded at a series of wavelengths spanning the steady-state emission spectrum of the dyes in steps of 10 nm. The fluorescence decays were fitted to a multi-exponential function via the reconvolution method using IBH DAS6 software. Three exponential components were necessary to obtain a satisfactory fit of the data. The purpose of the fit is to deconvolve the instrumental response from the data and should not be over-parameterized. The fitted decays together with the steady-state emission spectrum were used for the reconstruction of time-resolved emission spectra (TRES) by a spectral reconstruction method [27]. The reconstruction routine was implemented in Matlab. The position of TRES maximum ν*(t)* and its full-width at half-maximum *FWHM(t)* were inspected. Two main parameters describing polarity and mobility of the probed system were derived from ν*(t)*: The total amount of fluorescence shift Δν and the relaxation time τ_R_. The total amount of fluorescence shift Δν reflects the polarity of the environment of the probe and is calculated as:

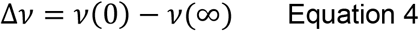

where ν*(0)* is the estimated position of TRES maximum before solvent relaxation [25,27], and ν(∞) is the position of the TRES at the fully relaxed state. The TDFS kinetics depends on the dynamics of the polar moieties in the vicinity of the probe and can be expressed as the integrated relaxation time τ_R_:

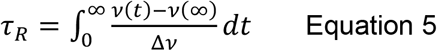

Intrinsic uncertainties for the TDFS parameters were 50 cm^−1^ and 0.05 ns for Δν and τ_R_, respectively.

### Molecular Dynamics Simulations

To construct the topologies of the fluorescent probes, we first performed the *ab initio* geometry optimization of the florescent fragments capped by the methyl groups. The B3LYP functional [29] with the 6-31G+(d, p) basis set [30–33] was used within the Gaussian 09 program [34]. Geometry optimization was performed for both the ground and first excited states of each fluorescent probe. To find the excited state geometries the TD-DFT scheme within the Gaussian 09 was employed. Optimized structures were all local minima as demonstrated by the frequency analysis. The optimized geometries of the first excited and ground states for most of the fluorescent probes did not differ substantially. However, in the case of Pro12A, a big difference in the molecular geometry of the ground and excited states was observed. The excited state geometries have been used in further studies. Subsequently, atomic charges were calculated using the Merz-Kollman-Singh ESP fit for the optimized structures [35]. After removing the terminal methyl groups, the fluorescent probes were attached to the lipid chains and GROMACS topologies were built.

In this study, we performed atomistic molecular dynamics (MD) simulations of single-component lipid bilayers composed of POPC, and mixed bilayers composed of POPC and 20 mol% of cholesterol. Bilayers were comprised of 200 lipid molecules (POPC and cholesterol) and eight fluorescently labelled compounds (4 mol%). The detailed composition of models used in this study is given in Supplementary Table 1. The initial configurations of all models were prepared with the CHARM-GUI server [36], and the GROMACS 2021 software package was used to perform MD simulations [37]. All models were simulated with three independently constructed replicates each 1 µs long, including 200 ns of equilibration. The all-atom OPLS force field was used to parameterize lipids and fluorescent labels [38,39]. Additional parameters necessary to describe lipids molecules were taken from the literature [40,41]. A topology for POPC was taken from ref [42], and the topology for cholesterol molecules was taken from ref [43]. For water, the TIP3P model was used [44].

Periodic boundary conditions were applied in all three directions. Simulations were performed under constant temperature (300 K) and in constant pressure (1 atm). Temperature and pressure were controlled with the Nose-Hoover thermostat [45] and Parrinello−Rahman barostat [46], respectively. A coupling constant for temperature was 0.4 ps. The temperature of bilayers (POPC, cholesterol, and probes) and water (water and counterions) were controlled independently. The coupling constant for pressure control was 1 ps. A semi-isotropic scheme of pressure coupling was applied (pressure in the membrane plane was controlled separately from pressure along the bilayer normal). Covalent bonds in lipids and fluorescence probes were restrained using the LINCS algorithm [47], which allows for the 2 fs time step. For the bonding interactions in water molecules, the SETTLE algorithm was employed [48]. The particle mesh Ewald (PME) algorithm was used for the treatment of long-range electrostatic interactions, and the cutoff used for the Lennard-Jones potential was chosen to be 1 nm.

## Results and Discussion

### Sensitivity of Pro12A, NR12S and NR12A to membrane phases

Independent studies have shown that NR12S, NR12A and Pro12A are able to report on the degree of lipid packing, and can therefore distinguish membrane fluidity in the plasma membrane [14–16]. Heterogeneity in membrane order can be reconstituted in phase-separated model membranes. Ordered phases are characterized by saturated lipids and high cholesterol content, whereas disordered phases comprise mostly unsaturated lipids [49]. Here we compared the ability of these probes to discriminate between ordered and disordered phases using two different systems, namely giant unilamellar vesicles (GUVs) (obtained by mixing sphingomyelin (SM), 1,2-dioleoyl-sn-glycero-3-phosphocholine (Δ9*cis* DOPC, 18:1/18:1 PC) and cholesterol in a 2:2:1 ratio) and giant plasma membrane vesicles (GPMVs) derived from U2OS cells.

With the use of these probes (structures depicted in Figure 1A) in advanced imaging methods such as spectral imaging, we can study the emission shift as a response to the environment in ∼ 9 nm wavelength intervals. The spectral images obtained for Pro12A, NR12S and NR12A are shown in Figure 1B,C and D, respectively. The normalized intensity of ordered and disordered phases in each of the channels shows the spectral shift of the individual probes in phase separated GUVs and GPMVs (Figure 1E-G). For all three fluorescent probes a clear shift in emission towards red wavelengths from ordered to disordered phase was observed in phase separated GUVs, with the biggest shift exhibited by Pro12A and the smallest by NR12S. In phase separated GPMVs the red shift was less visible than in GUVs as expected due to the compositional complexity [24,50,51], especially for NR12A.

To quantify the spectral shift of the probes in response to the environment, we calculated the generalized polarization (GP) value, which reflects membrane order (i.e., higher GP value corresponds to higher membrane order) [18]. For calculation of GP, two reference wavelengths (one in the blue-shifted and one in the red-shifted spectral region) must be chosen. We picked those two wavelengths in which the differences in intensity of ordered and disordered phases were the highest and led to the largest GP range. The respective wavelengths chosen for GP value calculation for the probes are 423 and 494 nm for Pro12A, 557 and 664 nm for NR12S, and 583 and 673 nm for NR12A (indicated by the coloured boxes in Figure 1B-D and dashed lines in Figure 1E-G). Of note, optimization for best GP contrast might come at a cost of noise (hence larger spread of data) due to low signal at the wavelengths chosen. When comparing different smart probes, absolute GP values are not informative since they are probe dependent. Therefore, we compared the performance of the three probes by examining the difference between GP values obtained from ordered and disordered phases within the same phase-separated vesicle. As shown from the colour-coded GP maps and linearized GP profiles along the vesicle (Figure 1H-J), all three probes are responsive to lipid packing in the membrane. However, NR12A shows a smaller spectral red-shift in disordered phases in GPMVs, which is especially apparent in the linearized GP profile along the phase-separated vesicle (Figure 1J).

As expected, for all probes the ΔGP is smaller in phase separated GPMVs compared to GUVs as these model membranes are derived from cells with a large compositional diversity (Figure 1K-M, Supplementary Figure 1A-C). Pro12A exhibits large GP-ranges with a ΔGP of 0.76 ± 0.16 (mean of three independent replicates ± standard deviation) in GUVs and a ΔGP of 0.63 ± 0.10 in GPMVs and can therefore separate ordered and disordered phases very well (Figure 1K), confirming a previous study [14]. NR12S has the smallest ΔGP of 0.59 ± 0.12 in GUVs, but dissects ordered and disordered phases well in GPMVs with a ΔGP of 0.45 ± 0.14 (Figure 1L). NR12A is superior to NR12S in differentiating the phases in GUVs with a ΔGP of 0.87 ± 0.15 as previously reported (Figure 1M) [15]. However, NR12A exhibits a smaller range in GPMVs with a ΔGP of 0.27 ± 0.05.

### Pro12A, NR12S and NR12A exhibit different sensitivities to saturation, double bond position and configuration, headgroup and cholesterol content

As examined above, the three probes distinguish between lipid phases in synthetic and cell-derived membranes; however, they show differences in ΔGP for ordered to disordered phases depending on the model membrane system. This indicates that the probes may differ in their sensitivity to different biophysical properties in their immediate membrane environment. To pinpoint individual sensitivities of Pro12A, NR12S, and NR12A to different membrane properties, we examined their spectral properties in well-defined membrane compositions. Specifically, we calculated GP values as a function of *i)* the degree of saturation, *ii)* position and configuration (*cis* vs. *trans*) of the double bond, *iii)* phospholipid headgroup and *iv)* cholesterol content (Figure 2A).

**Figure 2:**
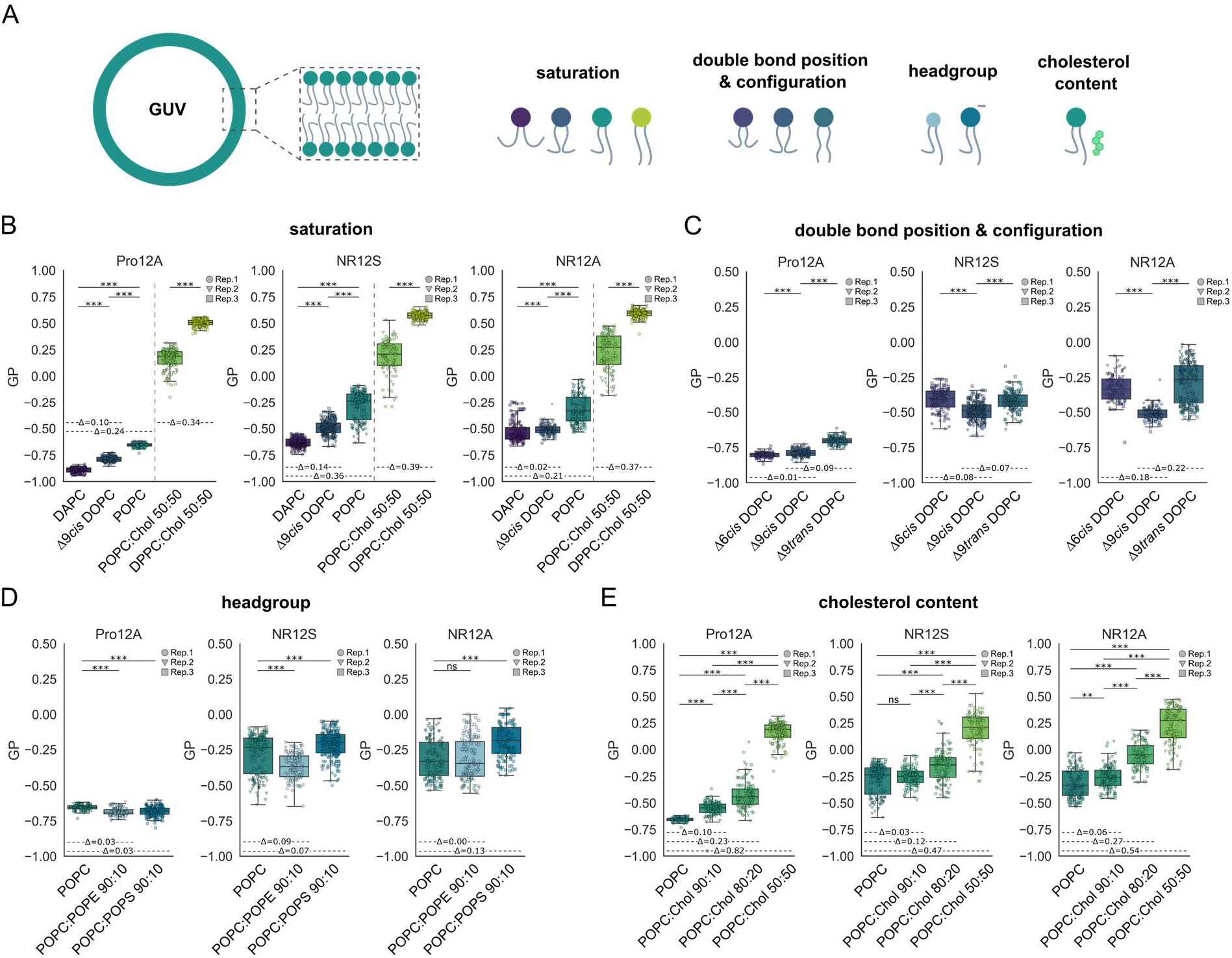
Sensitivities of Pro12A, NR12S and NR12A to changes in membrane composition. Spectral imaging and subsequent GP calculation of the probes in GUVs of varying lipid composition. A| Schematic overview. B| Saturation. GP of the probes in membranes with increasing degree of saturation. C| Double bond position and configuration. GP of the probes in DOPC membranes with Δ6 vs. Δ9 position and *cis* vs. *trans* configuration. D| Headgroup. GP of the probes in POPC membranes with 10 mol% of either POPS or POPE. E| Cholesterol content. GP of the probes in POPC membranes with increasing amount of cholesterol. Colours indicate the different lipid compositions. Data of three independent experiments is shown and replicates are indicated by data point shape. Values on the horizontal dashed lines correspond to the ΔGP values calculated from the mean. Kruskal Wallis test with post-hoc Mann-Whitney U test were performed for testing of significance (α=0.05).

#### Saturation

The degree of saturation of lipids impacts membrane properties [49]. Unsaturated lipids possess one or more double bonds, which are usually of *cis* conformation and introduce kinks in an acyl chain. This increases membrane fluidity by hindering dense lipid packing, with a consequent increase in lipid mobility and membrane hydration [49]. High uptake of saturated fatty acids is linked to obesity and diabetes, causing increased membrane saturation in e.g. erythrocytes [52], as well as ER stress and toxicity in hepatocytes [53]. In cancer cells, increased saturation stabilizes membrane domains resulting in increased signalling, and potentially chemoresistance [54]. Therefore, it is crucial to have a straightforward way to assess the lipid saturation in various biological systems using smart probes.

To examine how the probes respond to the degree of saturation, we prepared GUVs using 1,2-diarachidonoyl-sn-glycero-3-phosphocholine (DAPC, 20:4/20:4 PC) as polyunsaturated lipid, 1,2-dioleoyl-sn-glycero-3-phosphocholine (Δ9*cis* DOPC, 18:1/18:1 PC) as double-unsaturated lipid and 1-palmitoyl-2-oleoyl-glycero-3-phosphocholine (POPC, 16:0-18:1 PC) as mono-unsaturated lipid. We also wanted to test the fully saturated lipid 1,2-dipalmitoyl-sn-glycero-3-phosphocholine (DPPC, 16:0/16:0 PC); however, it forms GUVs only in the presence of additional cholesterol. Therefore, we used POPC and DPPC with 50 mol% cholesterol (POPC:Chol vs. DPPC:Chol) to compare between monounsaturated and fully saturated lipids in the presence of the same amount of cholesterol. All probes can reliably differentiate between the different degrees of lipid saturation in their environment and display an increasing GP with increasing degree of saturation (Figure 2B), yet to a different extend. Among the three probes, NR12S is the most sensitive to saturation (ΔGP_DAPC/DOPC_ = 0.14; ΔGP_DAPC/POPC_ = 0.36; ΔGP_POPC:Chol/DPPC:Chol_ = 0.39). NR12A is good at distinguishing between monounsaturated and saturated lipids, but it is the least powerful of the three probes at differentiating among the unsaturated lipids. Although Pro12A shows the overall largest ΔGP range, the individual GP differences are smaller compared to NR12S. In general, all probes differentiate monounsaturated and saturated lipids much better than between unsaturated lipids.

#### Configuration

Similar to the degree of saturation, position or configuration (*cis* vs. *trans*) of the double bond impacts membrane properties [6]. Increased uptake of *trans*-unsaturated fatty acids as well as a high ratio of omega-6 to omega-3 polyunsaturated fatty acids alter the plasma membrane composition and are thereby linked to risk for diabetes, obesity, cancer, systemic inflammation and cardiovascular disease [55–57]. Therefore, we examined whether the probes sense variations in position or configuration of the double bond. To this end, we prepared GUVs from lipids only differing in double bond position and configuration, namely 1,2-dipetroselenoyl-sn-glycero-3-phosphocholine (Δ6*cis* DOPC, 18:1/18:1 PC), Δ9*cis* DOPC, and 1,2-dielaidoyl-sn-glycero-3-phosphocholine (Δ9*trans* DOPC, 18:1/18:1 PC) (Figure 2C). NR12A is more sensitive to double bond position (ΔGP_Δ6*cis/*Δ9*cis*_ = 0.18) compared to NR12S (ΔGP_Δ6*cis/*Δ9*cis*_ = 0.08), whereas Pro12A appears to be insensitive (ΔGP_Δ6*cis/*Δ9*cis*_ = 0.01). Theoretically, the position of the double bond determines where the kink of the acyl chain is located and how deep water molecules can penetrate the membrane, which explains the decrease in GP from Δ6*cis* DOPC to Δ9*cis* DOPC for NR12A and NR12S [58]. Pro12A is not able to sense this difference in water penetration, which indicates that it might adopt a distinct location from that of NR12A and NR12S (discussed later).

In theory, a *trans* configuration of the double bond straightens the acyl chain, therefore allowing the lipids to pack more densely and causing an increase in GP [58,59]. Indeed, all three probes show an increase in GP for Δ9t*rans* DOPC compared to Δ9*cis* DOPC, with NR12A being the most sensitive (ΔGP_Δ9*cis/*Δ9*trans*_ = 0.22), followed by Pro12A (ΔGP_Δ9*cis/*Δ9*trans*_ = 0.09) and NR12S (ΔGP_Δ9*cis/*Δ9*trans*_ = 0.07).

#### Headgroup

The headgroups of phospholipids, which can vary in size and charge, not only drive certain cellular processes by lipid-protein interaction, but also structurally define membrane properties [49]. Phospholipids are asymmetrically distributed in the membrane bilayer with phosphatidylcholine (PC) and sphingolipids concentrated in the outer leaflet, whereas phosphatidylserine (PS) and phosphatidylethanolamine (PE) mostly reside in the inner leaflet [4,60]. Accounting for 10-15 % of plasma membrane lipids, negatively charged PS serves as a signalling platform [61] and is involved in establishing membrane potential [62]. Translocation of PS to the outer membrane leaflet in apoptosis, necroptosis and pyroptosis [60] can trigger engulfment by macrophages [63], the coagulation cascade in platelet activation [62], and might have implications in calcium-regulated exocytosis [64]. PE is a zwitterionic lipid with a small headgroup to acyl chain ratio resulting in a conical shape that induces negative curvature and packing defects, increasing the membrane fluidity [49]. At the plasma membrane, PE plays an important role in fusion and fission events and membrane topology, enhances the integration of proteins into the bilayer [65] and its translocation to the outer leaflet is essential for cell-cell fusion [66]. Thus, we asked whether the translocation of PS and PE to the outer bilayer leaflet during above-mentioned events could be studied using our smart probes residing in the outer leaflet. In other words, how do our smart probes react to different headgroups?

To investigate the influence of phospholipid headgroup on the spectral shift of the probes, we generated GUVs comprising either pure POPC or mixed with 10 mol% of 1-palmitoyl-2-oleoyl-sn-glycero-3-phospho-L-serine (POPS, 16:0/18:1 PS) (confirmed by Annexin V-Alexa Fluor 647 staining, Supplementary Figure 2) or 1-palmitoyl-2-oleoyl-sn-glycero-3-phosphoethanolamine (POPE, 16:0/18:1 PE). Both POPS and POPE share the same acyl chain moiety with POPC. Therefore, we can investigate whether the probes are sensitive to headgroup charge, using POPS, and headgroup size or geometry, using POPE.

The three probes exhibit different sensitivities to POPE and POPS (Figure 2D). Overall, Pro12A shows the least sensitivity to headgroup charge (POPS) and size or geometry (POPE), (ΔGP = 0.03 for both compared to pure POPC). NR12S is moderately sensitive to headgroup charge (ΔGP_POPC/POPS_ = 0.07) and size or geometry (ΔGP_POPC/POPE_ = 0.09). While NR12A exhibits the largest sensitivity to headgroup charge (ΔGP _POPC/POPS_ = 0.13) among the probes, it cannot distinguish headgroup size or geometry at all. The differences in ΔGP as a function of headgroup charge and size or geometry, may arise due to different orientations and locations of the probes in the membrane influencing their electronic relaxation process, which we will address in later sections.

#### Cholesterol

In addition to variations in phospholipid structure, cholesterol content has a large impact on membrane properties and fluidity and is thereby crucial for many cellular processes occurring at the plasma membrane [49]. In fact, increased cholesterol levels are linked to metabolic diseases such as hypercholesteremia and atherosclerosis [53]. Cancer cells often exhibit increased cholesterol uptake, synthesis and storage. Their plasma membrane cholesterol content varies: high cholesterol drives membrane domain-associated signalling and drug resistance [54], whereas low cholesterol increases membrane fluidity crucial for cell mobility and infiltration of other tissues in metastasis [67]. To assess the usability of our probes for such biological questions, we tested how they react to different cholesterol levels in membranes.

We investigated the sensitivity of Pro12A, NR12A and NR12S to cholesterol content by examining their spectral shift in POPC GUVs containing increasing amounts of cholesterol (Figure 2E). All probes can reliably report on cholesterol content, with Pro12A being the most sensitive (mean ΔGP_POPC/POPCChol50_ = 0.82; ΔGP_POPC/POPCChol10_ = 0.10), followed by NR12A (ΔGP_POPC/POPCChol50_ = 0.54; ΔGP_POPC/POPCChol10_ = 0.06) and NR12S, which however fails to sense low cholesterol content (ΔGP_POPC/POPCChol50_ = 0.47; ΔGP_POPC/POPCChol10_ = 0.03, not significant). Therefore, small changes in cholesterol content might not be probed by NR12S.

#### Summary of different sensitivities

Taken together, Pro12A exhibits the biggest change in GP (ΔGP = 1.40) from the most disordered (DAPC) to most ordered (DPPC:Chol 50:50) membrane composition investigated, followed by NR12S (ΔGP = 1.21) and NR12A (ΔGP = 1.12). Pro12A is highly sensitive to cholesterol content as well as lipid saturation and can distinguish the configuration of the DOPC double bond. Nonetheless, it is almost insensitive to phospholipid headgroups such as POPS and POPE, and double bond position in DOPC, which can be an advantage or disadvantage depending on the biological question. NR12S is superior at differentiating lipids of varying saturation and is the only probe that can differentiate among all headgroups investigated. It can also distinguish both double bond position as well as configuration in DOPC but is the least sensitive to cholesterol content. Finally, NR12A is superior at differentiating double bond position and configuration in DOPC and senses the presence of POPS. NR12A is sensitive to cholesterol content and lipid saturation but performs better with more saturated than poly-unsaturated lipids. Notably, the GP values of Pro12A are the least widely distributed in all lipid compositions examined compared to NR12S and NR12A, which makes it easier to compare different lipid bilayers. All these differences are likely due to the probe localization and orientation in the membrane, which we address next.

### Atomistic molecular dynamics simulations reveal distinct probe locations in the bilayer

The probe sensitivity to membrane properties is influenced by its orientation and position within the membrane bilayer [68,69]. Therefore, we performed *in silico* atomistic MD simulations on each individual probe to gain more insight regarding its location and orientation [70]. Specifically, we analysed two different scenarios: the probes in *i)* POPC and *ii)* POPC:Chol lipid bilayers. For a summary of the MD simulation analysis of all probes see Supplementary Tables 1-3. The position and orientation of the probes in these bilayers are depicted in snapshots (Figure 3A). To examine the orientation of the probes within the membrane bilayer, we determined the tilt angle distributions between the membrane normal and the long axes of the fluorescent moieties (Figure 3B). If the tilt angle is 0 °, then the probe is oriented along the membrane normal direction, pointing towards the centre of the membrane. If the tilt angle is 90 °, then the probe is oriented along the membrane surface. In POPC membranes, Pro12A and NR12S show an orientation that is largely parallel to the bilayer normal with average tilt angles of 38.5 ° and 33 °, respectively (Supplementary Table 2). Interestingly, NR12S exhibits two maxima in the tilt angle distribution at 23 ° and 87 °, indicating that a small fraction of NR12S molecules is oriented parallel to the membrane surface. Bimodal distribution of solvatochromic dyes in lipid bilayers was noticed previously and assigned to capacity of these dyes to form H-bonds with water leading to shallow located probe population [71]. NR12A has an average tilt angle of 71 ° and is therefore oriented largely parallel to the membrane surface. The presence of 20 mol% cholesterol in the lipid bilayer slightly reduces the average tilt angle by ∼ 5 °, ∼2 ° and ∼1 ° for NR12S, NR12A and Pro12A, respectively. This is in line with cholesterol’s ability to increase lipid packing [5], squeezing the lipids more closely together and thereby decreasing the tilt angle of the probes. We also mapped (via partial density profiles) the distribution of specific atoms, selected from both the probe and the lipids, along the lipid bilayer to estimate the location of the probes within the membranes (Figure 3C). Reflecting the parallel orientation, Pro12A and NR12S are distributed along the whole bilayer normal (wide partial density profiles of the probe), with NR12S locating closest to the water phase (see Supplementary Table 3 for maxima of partial density profiles). Interestingly, the bimodality in the tilt angle distribution of NR12S is reflected in the wide partial density profile of the probe. In contrast, NR12A exhibits a more defined position in the hydrocarbon core of the bilayer between the POPC double bond and the membrane-water interface, due to its perpendicular orientation to the bilayer normal (narrower partial density profiles of the probe).

**Figure 3:**
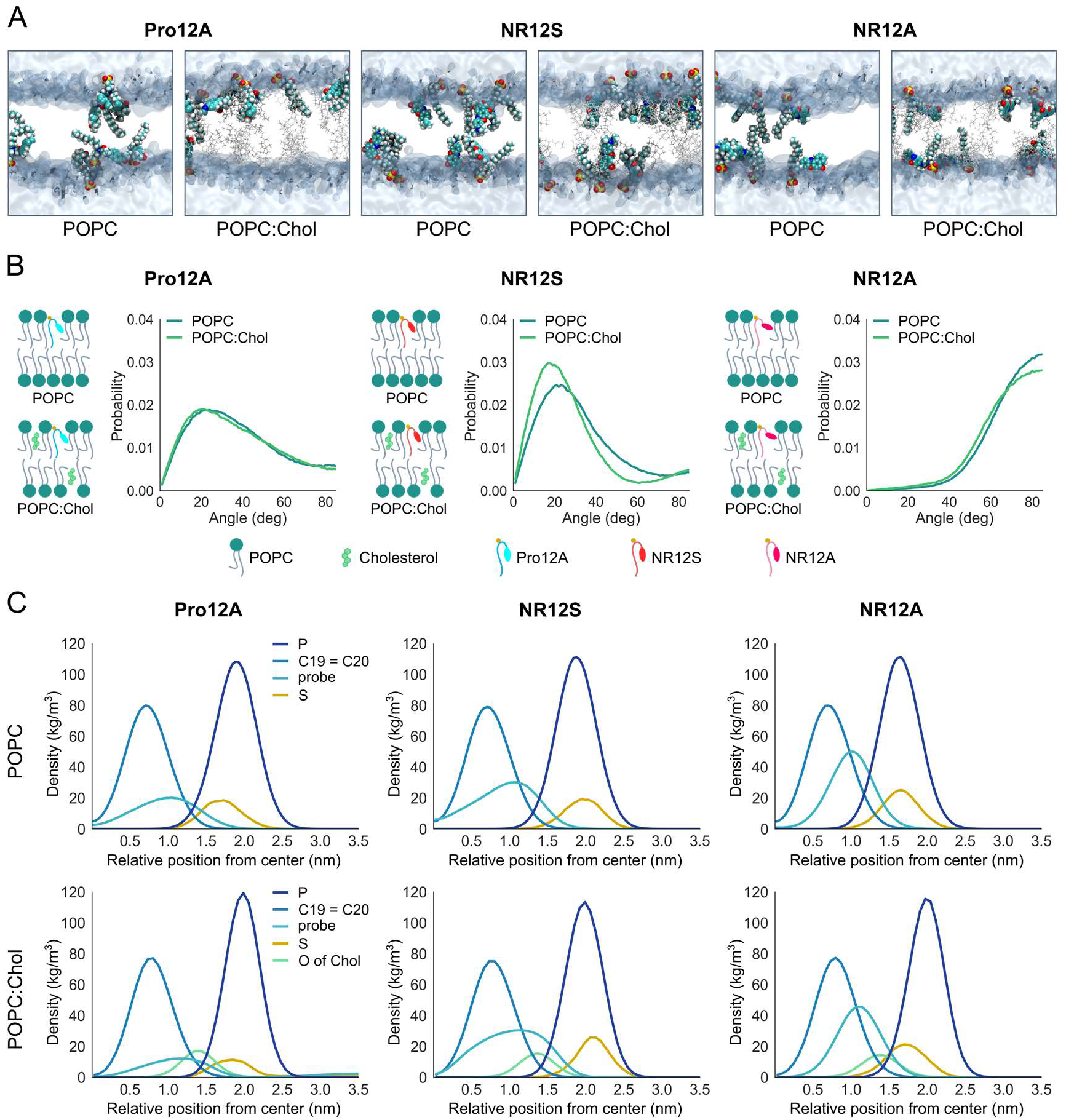
MD simulations of Pro12A, NR12S and NR12A disclose variations in probe orientation and location within the membrane. MD simulations were performed for membranes described in Supplementary Table 1. A| Visualizations of the probes’ orientation and location in POPC and POPC:Chol (80:20) membranes. Probe, cholesterol and water molecules are shown in the VDW, licorice, and surface representation, respectively. POPC is not shown for clarity. B| Tilt angle distributions of the three probes in POPC and POPC:Chol (80:20) membranes. C| Partial density profiles of the probes and selected atoms of the lipids to determine the location of Pro12A, NR12S and NR12A in POPC and POPC:Chol (80:20) membranes.

In POPC bilayers cholesterol is located deep in the membrane. The hydrocarbon tail localizes at the centre of the bilayer and the OH-group faces towards the headgroups of the surrounding lipids (upright position) [72]. This is confirmed by our MD simulation data, which locates the cholesterol OH-group at ∼1.4 nm within the ∼3 nm thick bilayer. The partial density profiles of the three probes show proximity to cholesterol, supporting their sensitivity to different cholesterol amounts in the membrane. Addition of 20 mol% cholesterol to the POPC membrane slightly shifts the partial density profiles for the fluorescent moiety and sulphur atom of all the probes outwards, closer to the water phase (∼ 0.2 nm). This is likely a result of the increase in lipid packing due to cholesterol. The shoulder of the partial density profile of NR12S’ fluorescent moiety is even more pronounced in POPC:Chol, indicating that NR12S can assume multiple locations along the whole bilayer and a small fraction of the probe molecules adopts an orientation perpendicular to the bilayer normal. The dual location of NR12S is in line with previous molecular simulation studies [73] and with observed poor two-photon light polarization effects for NR12S in ordered phases of GUVs [16]. This overall less defined location of NR12S in the membrane, resulting in different immediate environments of the probe, might be the reason why NR12S cannot resolve low cholesterol content (10 mol%, as observed in GP analysis above).

Although POPC and POPC:Chol bilayers exhibit different membrane properties, the orientation and location of the probes remain relatively unaffected. However, each individual probe has its own specific localization within the membrane due to their molecular structure. This distinct localization of the probes explains their distinct sensitivities to different membrane properties.

### TDFS

The observed variations in orientation and position of the probes indicate that they are exposed to different immediate environments even within membranes of the exact same lipid composition. The immediate environment affects the probes’ dipolar relaxation process, which is the underlying principle of the emission shift, i.e., the emission shift to longer wavelengths depends on the polarity and viscosity of the solvent [74]. The dipolar relaxation process can be examined by the TDFS method, which provides detailed information on the fluidity as well as polarity level of the hydrated phospholipids in the lipid bilayer area surrounding the probe [75,76]. Using TDFS, it has previously been shown that Laurdan has a single relaxation process, whereas di-4-ANEPPDHQ has multiple processes that are likely caused by its electrochromic sensitivities [12]. That study showed that probes might have different dipolar relaxation dynamics, and one cannot assume that each environment-sensitive probe reports on the same aspect of the membrane. This is particularly crucial when applying such probes in a complex cellular context. Thus, we examined the dipolar relaxation process of our probes via TDFS.

In TDFS, solvent relaxation is commonly characterized by the relaxation time τ_R_ and overall dynamic Stokes shift Δν. τ_R_ is related to the time taken by the probe’s environment to reorient due to the new energetic state of the probe, therefore τ_R_ is a function of the *fluidity* of the probe environment. In other words, in more fluid bilayers, τ_R_ is shorter because it takes shorter time for the hydrated lipids in the environment to move around the probe. Δν is the difference between the non-relaxed excited state and fully-relaxed electronic excited states of the probe once the probe is excited and it is a function of *polarity* [76]. In other words, in more polar environments, Δν is higher.

We investigated the relaxation processes of the probes in large unilamellar vesicles (LUVs) of different lipid compositions (Figure 4A,B, Supplementary Figure 3, summary of the TDFS analysis of all probes can be found in Supplementary Tables 4-6). For Pro12A, τ_R_ becomes longer with increasing membrane order (i.e., increasing GP), indicating that the probe can differentiate well the degree of fluidity among hydrated poly-, mono-unsaturated and saturated lipids as well as cholesterol content (Figure 4A). For POPC:Chol 50:50 and more so for DPPC:Chol, the relaxation time is underestimated as the relaxation process is longer than the fluorescence lifetime of Pro12A (see Supplementary Table 4 and Supplementary discussion). Of note, the time profiles full width half maxima (FWHM) of the time resolved emission spectra (TRES) of Pro12A (see Supplementary Figure 3C,D) behave exemplarily as anticipated for typical solvent relaxation probes (see Supplementary discussion). In fact, there is a strong experimental evidence suggesting that Prodan (and derivatives) is an ideal solvent relaxation chromophore [77,78].

**Figure 4:**
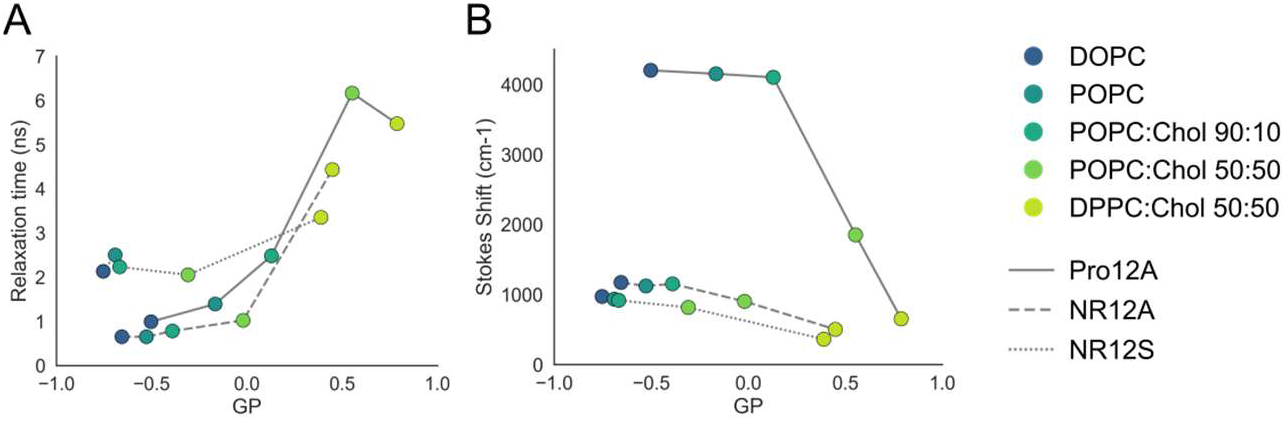
TDFS analysis of Pro12A, NR12S and NR12A reveals variations in relaxation behaviour in different lipid environments. TDFS analysis of the probes was performed in LUVs of varying lipid composition at 23 °C. Summary of the TDFS analysis of all probes can be found in Supplementary Tables 4-6. A| Relationship of relaxation time and GP-value for all probes. B| Relationship of Stokes shift and GP-value for all probes. The line style indicates the probe. Colours indicate the different lipid compositions.

For NR12A the increase of τ_R_ with higher membrane order modulated by increasing cholesterol content is less pronounced and τ_R_ surpasses 1 ns only in the more ordered bilayers (POPC:Chol 50:50 and DPPC:Chol) (Figure 4A, Supplementary Figure 3A,G,H, Supplementary Table 6). This suggests that the dipolar relaxation of NR12A is not as sensitive to cholesterol content as Pro12A which is in line with experimental findings (see Figure 2). Similarly, for NR12S there is a slight increase of τ_R_ with increasing GP for most compositions (Figure 4A, Supplementary Figure 3A,E,F, Supplementary Table 5), and τ_R_ increases significantly for DPPC:Chol suggesting that the probe is less sensitive to membrane fluidity changes caused by varying cholesterol content, but more sensitive when saturated lipids are present.

The Stokes shift parameter of TDFS, Δν, is determined from the position of the TRES maximum and reports on the hydration level in the probes’ vicinity (i.e., polarity). The higher the hydration in the vicinity of the probe the larger the Stokes Shift [76]. The Δν of Pro12A remains constant in DOPC, POPC and POPC:Chol 90:10 (Figure 4B, Supplementary Figure 3B, Supplementary Table 4), implying the probe cannot sense a difference in membrane polarity upon switch from poly-unsaturated to mono-unsaturated lipids or low cholesterol content. In the case for NR12S and NR12A, Δν is difficult to interpret, nevertheless the observed decrease in Δν for POPC:Chol and in DPPC:Chol suggests a decrease in polarity in the bilayer upon swap from mono-unsaturated to fully saturated lipids or high cholesterol content (Figure 4B, Supplementary Figure 3B, Supplementary Tables 5,6 and Supplementary discussion). The probe’s sensitivity to polarity is also reflected in the increase of the GP value [79].

In summary, the TDFS data shows that Pro12A is sensing the fluidity (estimated from the relaxation time) of the hydrated lipid segments in its immediate environment rather than the hydration level of the membrane itself (estimated from the Stokes shift). Therefore, one can conclude that the GP parameter for this dye is mainly the reflection of the fluidity of the environment of the probe. For NR12A and NR12S, the GP value is affected by both the membrane’s fluidity and hydration. However, for these probes, an internal photophysical process (such as twisting in the excited state suggested for Nile Red [80,81]) might contribute in addition to the dipolar relaxation and the TDFS data interpretation is not as straightforward as in the case of Pro12A. These processes might partially account for the large standard deviations of data obtained using NR12A and NR12S, in contrast to the data obtained using Pro12A with narrower spread (Figure 2). Finally, this intricacy of NR12A and NR12S confirms that these probes are sensitive to different aspects of membranes compared to Pro12A as shown experimentally (Figure 2).

## Conclusions

Environment-sensitive fluorescent probes have greatly contributed to our understanding of cellular membranes. They enable the investigation of biophysical properties, phase-separation and lipid domain formation in membranes. After Laurdan was introduced [10], various probes have been developed in the last two decades with improved partitioning into the membrane, selectivity for the outer membrane leaflet, increased brightness and photostability for advanced imaging, and selectivity for organelle-specific membranes [8,15,82,83]. These probes are assumed to report on membrane order in similar ways. In this study, we examined the membrane probes Pro12A, NR12S and NR12A in model membranes of defined lipid compositions and revealed their different sensitivities to the degree of lipid saturation, double bond position and configuration, chemical details in phospholipid headgroup and membrane cholesterol content. Knowledge of these sensitivities is highly beneficial for application of these probes in different biological contexts. The present study serves as a guide on when to use each probe and what their advantages and disadvantages can be. Additional TDFS analyses and atomistic MD simulations of the probes contribute to the understanding of how these sensitivities arise from molecular details by providing insight into the relaxation process as well as information on orientation and position of the probes within the bilayer, respectively.

As revealed by this work, the probe Pro12A is the most sensitive to cholesterol content in the membrane, but also performs well at differentiating between the degree of saturation and configuration of the double bond. Moreover, data obtained with Pro12A has a significantly smaller standard deviation, giving it power to distinguish even smaller differences. NR12S is superior at reporting the degree of lipid saturation and changes in phospholipid headgroup structure, but also responds well to double bond position and configuration. NR12A distinguishes the position and configuration of the double bond best and recognizes changes in cholesterol content and degree of saturation.

The probes’ individual sensitivities can direct their application in investigating the plasma membrane in various biological contexts. Therefore, instead of choosing one probe over another one, we recommend using the one best-suited for the hypothesis in question or to apply all of them to gain a full picture of the different membrane properties.

It should be noted that the probes’ sensitivities were examined in membrane model systems of certain defined lipid composition. First, it might also be of interest to investigate the influence of other lipids such as sphingomyelin (particularly in the presence of cholesterol), plasmalogens and other sterols. Moreover, cellular membranes exhibit much higher lipid heterogeneity and additionally comprise membrane-embedded proteins, which can alter membrane properties due to protein-protein or lipid-protein interactions. Therefore, live cells represent a much more complex system and further work on these probes is needed.

## Acknowledgement

We thank SciLifeLab Advanced Light Microscopy facility and National Microscopy Infrastructure (VR-RFI 2016-00968) for their support on imaging. We would like to acknowledge Mgr. Eduard Jesko and Tatjana Šafarik for their contribution to TDFS experiments in the early stages of the project. ES is supported by Swedish Research Council Starting Grant (2020-02682), SciLifeLab National COVID-19 Research Program financed by the Knut and Alice Wallenberg Foundation, Cancer Research KI, Human Frontier Science Program. FR is supported by Karolinska Institutet KID grant. MH, JS and MA acknowledge GACR grant 19-26854X. DID was supported by a fellowship from the Ministère de la Recherche (France). IV acknowledges support granted by the Academy of Finland (project no. 331349, 346135), Human Frontier Science Programme (RGP0059/2019), Sigrid Juselius Foundation, the Helsinki Institute for Life Science (HiLIFE), and Cancer Foundation Finland. We also acknowledge CSC - IT Center for Science (Espoo, Finland) for providing computing resources.

