## Supplementary Information for "Dissecting the mechanisms of environment sensitivity of smart probes for quantitative assessment of membrane properties"

### Deceased

#### Supplementary Tables

**Supplementary Table 1: Composition of model bilayers used in MD simulations, given as numbers of molecules or ions.**

| POPC | Chol | probe | K+ | water |
|---|---|---|---|---|
| 200 | - | Pro12A, 8 | 8 | 9100 |
| 160 | 40 | Pro12A, 8 | 8 | 9100 |
| 200 | - | NR12S, 8 | - | 9100 |
| 160 | 40 | NR12S, 8 | - | 9100 |
| 200 | - | NR12A, 8 | 8 | 9100 |
| 160 | 40 | NR12A, 8 | 8 | 9100 |

**Supplementary Table 2: Average and maximum tilt angles of the probes Pro12A, NR12S and NR12A as revealed by MD simulations.**

| | POPC | | POPC-Chol 80:20 | |
|---|---|---|---|---|
| | average (deg) | maximum (deg) | average (deg) | maximum (deg) |
| Pro12A | 38.5 | 24 | 37.9 | 21 |
| NR12S | 33 | 23 & 87 | 28.5 | 17 & 89 |
| NR12A | 71 | 86 | 69 | 86 |

**Supplementary Table 3: Maxima of the partial density profiles of sulphur and the fluorescence moiety of the probes Pro12A, NR12S and NR12A, given by analysis of MD simulations.**

| | POPC | | POPC cholesterol 80:20 | |
|---|---|---|---|---|
| | max. sulphur (nm) | max. probe (nm) | max. sulphur (nm) | max. probe (nm) |
| Pro12A | 1.73 | 1.05 | 1.85 | 1.15 |
| NR12S | 1.96 | 1.08 | 2.06 | 1.15 |
| NR12A | 1.65 | 1.01 | 1.70 | 1.07 |

**Supplementary Table 4: TDFS analysis of Pro12A in LUVs of varying lipid compositions at 23 °C and at 37 °C.**

**23 °C**

| lipid | Δν (cm-1) | $\tau_R$ (ns) | Peak time FWHM (ns) | observed relaxation (%) | GP value |
|---|---|---|---|---|---|
| DOPC | 4200 ± 100 | 0.99 ± 0.05 | 0.12 | 51 | -0.502 |
| POPC | 4150 ± 100 | 1.39 ± 0.05 | 0.32 | 62 | -0.166 |
| POPC/10Chol | 4100 ± 100 | 2.48 ± 0.05 | 1.39 | 74 | 0.129 |
| POPC/50Chol | 1850 ± 100* | 6.16* | >20 | 69 | 0.553 |
| DPPC/50Chol | 650 ± 100* | 5.47* | >20 | 86 | 0.788 |

**37 °C**

| lipid | Δν (cm-1) | $\tau_R$ (ns) | Peak time FWHM (ns) | observed relaxation (%) | GP value |
|---|---|---|---|---|---|
| DOPC | 4000 ± 50 | 0.59 ± 0.05 | <0.3 | 46 | -0.487 |
| POPC | 4100 ± 100 | 0.84 ± 0.05 | 0.14 | 56 | -0.342 |
| POPC/10Chol | 4100 ± 100 | 1.48 ± 0.05 | 0.48 | 61 | -0.117 |
| POPC/50Chol | 4000 ± 100 | 6.02 ± 0.05 | 5.56 | 87 | 0.358 |
| DPPC/50Chol | 750 ± 100* | >5.26* | >20 | 86 | 0.747 |

* Empirical parameters, final relaxed state cannot be fully probed as the relaxation process is not finished within the time period given by the fluorescence lifetime (average lifetime $\tau_{AV}$ is 4.9 ns and 6.3 ns for POPC/50Chol and DPPC/50Chol at 23 °C, respectively).

**Supplementary Table 5: TDFS analysis of NR12S in LUVs of varying lipid compositions at 23 °C and 37 °C.**

**23 °C**

| lipid | Δν (cm-1) | $\tau_R$ (ns) | Peak time FWHM (ns) | observed relaxation (%) | GP value |
|---|---|---|---|---|---|
| DOPC | 970 ± 100 | 2.13 ± 0.05 | -- | 77 | -0.753 |
| POPC | 931 ± 100 | 2.5 ± 0.05 | -- | 74 | -0.691 |
| POPC/10Chol | 915 ± 100 | 2.23 ± 0.05 | -- | 69 | -0.667 |
| POPC/50Chol | 813 ± 100 | 2.05 ± 0.05 | -- | 65 | -0.309 |
| DPPC/50Chol | 360 ± 100 | 3.35 ± 0.05 | -- | 79 | 0.389 |

**37 °C**

| lipid | Δν (cm-1) | $\tau_R$ (ns) | Peak time FWHM (ns) | observed relaxation (%) | GP value |
|---|---|---|---|---|---|
| DOPC | 1155 ± 50 | 2.02 ± 0.05 | -- | 70 | -0.791 |
| POPC | 1087 ± 100 | 1.88 ± 0.05 | -- | 68 | -0.732 |
| POPC/10Chol | 1111 ± 100 | 1.92 ± 0.05 | -- | 64 | -0.707 |
| POPC/50Chol | 997 ± 100 | 2± 0.05 | -- | 60 | -0.397 |
| DPPC/50Chol | 808 ± 100 | 2.27 ± 0.05 | -- | 54 | 0.297 |

**Supplementary Table 6: TDFS analysis of NR12A in LUVs of varying lipid compositions at 23 °C and 37 °C.**

**23 °C**

| lipid | Δν (cm-1) | $\tau_R$ (ns) | Peak time FWHM (ns) | observed relaxation (%) | GP value |
|---|---|---|---|---|---|
| DOPC | 1170 ± 100 | 0.65 ± 0.05 | -- | 37 | -0.655 |
| POPC | 1120 ± 100 | 0.65 ± 0.05 | -- | 49 | -0.527 |
| POPC/10Chol | 1150 ± 100 | 0.78 ± 0.05 | -- | 37 | -0.390 |
| POPC/50Chol | 900 ± 100 | 1.02 ± 0.05 | -- | 69 | -0.019 |
| DPPC/50Chol | 500 ± 100 | 4.43 ± 0.05 | -- | 100 | 0.449 |

**37 °C**

| lipid | Δν (cm-1) | $\tau_R$ (ns) | Peak time FWHM (ns) | observed relaxation (%) | GP value |
|---|---|---|---|---|---|
| DOPC | 1020 ± 50 | 0.74 ± 0.05 | -- | 45 | -0.647 |
| POPC | 1190 ± 100 | 0.71 ± 0.05 | -- | 44 | -0.539 |
| POPC/10Chol | 1160 ± 100 | 0.55 ± 0.05 | -- | 31 | -0.444 |
| POPC/50Chol | 1070 ± 100 | 0.91 ± 0.05 | -- | 39 | -0.127 |
| DPPC/50Chol | 760 ± 100 | 2.1 ± 0.05 | -- | 87 | 0.277 |

#### Supplementary discussion on TDFS results

The variation of the full width half maxima (FWHM) of the time resolved emission spectra (TRES) in time is a useful indicator of the extent of the observed relaxation process and its maximum can be used as an estimation of the average relaxation time (82). For the ideal TDFS probe and single dipolar relaxation processes, a FWHM time dependence reaches a single maximum approximately corresponding to the relaxation time $\tau_R$ (83). For the Pro12A dye a maximum of the FWHM over time can be observed in DOPC, POPC and POPC:Chol 90:10 but in POPC:Chol 50:50 and DPPC:Chol 50:50 membranes the relaxation process takes substantially longer than the fluorescence decay of the probe and a maximum in the FWHM profile is not visible (Supplementary Figure 3C). As the relaxation process is not finished within the time span given by the fluorescence lifetime of the dye, the Stokes shift parameter $\Delta\nu$ of Pro12A in POPC:Chol 50:50 and in DPPC:Chol 50:50 is ill estimated. Therefore, we cannot examine the complete relaxation process and the final solvent relaxed state cannot be probed rendering $\nu(\infty)$ an empirical parameter. The estimated values of the FWHM peak are shown in Supplementary Table 4. Increasing the temperature of the system increases its mobility and experiments performed at 37 °C showed faster relaxation dynamics as expected, and made it possible to observe a larger extent of the TDFS process and a FHWM peak for the POPC:Chol 50:50 sample. However this was not sufficient to do the same for the DPPC:Chol sample (Supplementary Figure 3D). This temperature dependent behaviour (also reflected in the GP results) further shows how the Pro12A dye is responsive to membrane mobility and that the dye is able to distinguish between the two more rigid samples of POPC:Chol 50:50 and DPPC:Chol (Supplementary Table 4, Supplementary Figure 3A,B).

For the case of the NR derived dyes, no apparent maxima in the FWHM profiles are detected at 23 and 37 °C (Supplementary Figure 3E-H). This behaviour anomalous for the classical TDFS lipid membrane probes, can be elucidated by a closer look into the Nile Red photophysics. Upon the excitation, a twisted internal charge transfer (TICT) takes place (53,54). This process competing with the solvent relaxation results in the time-dependent formation of various emitting species with different fluorescence properties being affected by the TICT propagation. This complex photophysical behaviour smears the FWHM time dependence abolishing the single maximum profile observed for Pro12A. In contrast to NR dyes, “Dan” (6-(dimethylamino)-2-acylnaphthalene) dyes, that Pro12A belongs to, have been proven to stay planar in the excited state (50,51), exhibiting solvation-dependent emission from a single excited-state entity.

#### Supplementary Figures

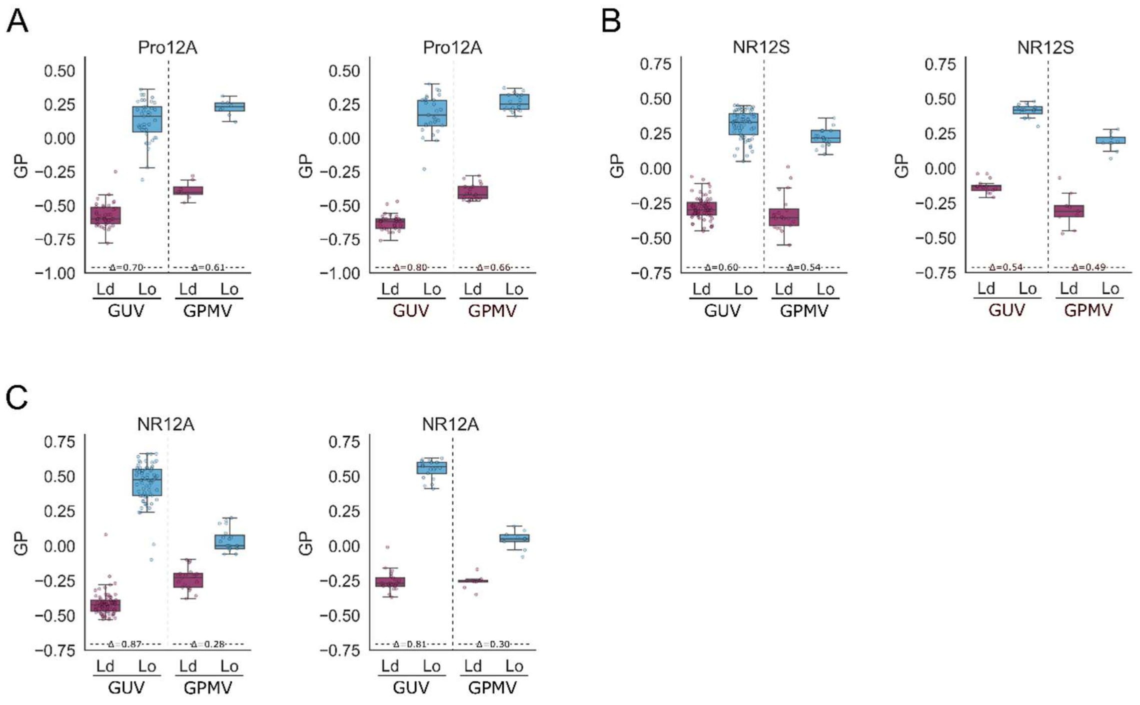


**Supplementary Figure 1: GP values of Pro12A, NR12S and NR12A in phase-separated GUVs and GPMVs**.

A-C| Calculated GP values of ordered and disordered phases in GUVs and GPMVs stained with Pro12A (A), NR12S (B) and NR12A (C). Each graph corresponds to another replicate.

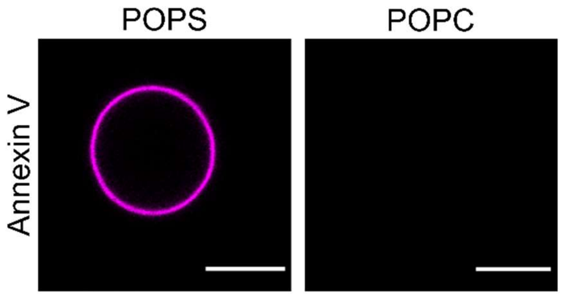


**Supplementary Figure 2: Annexin-V staining confirms integration of POPS into the membrane during GUV formation.** GUVs of POPC and 10 mol% POPS were performed with electroformation. Subsequent staining with Annexin-V 647 confirmed the successful integration of POPS into the membrane. Pure POPC GUVs were used as control. Scale bar = 10 µm.

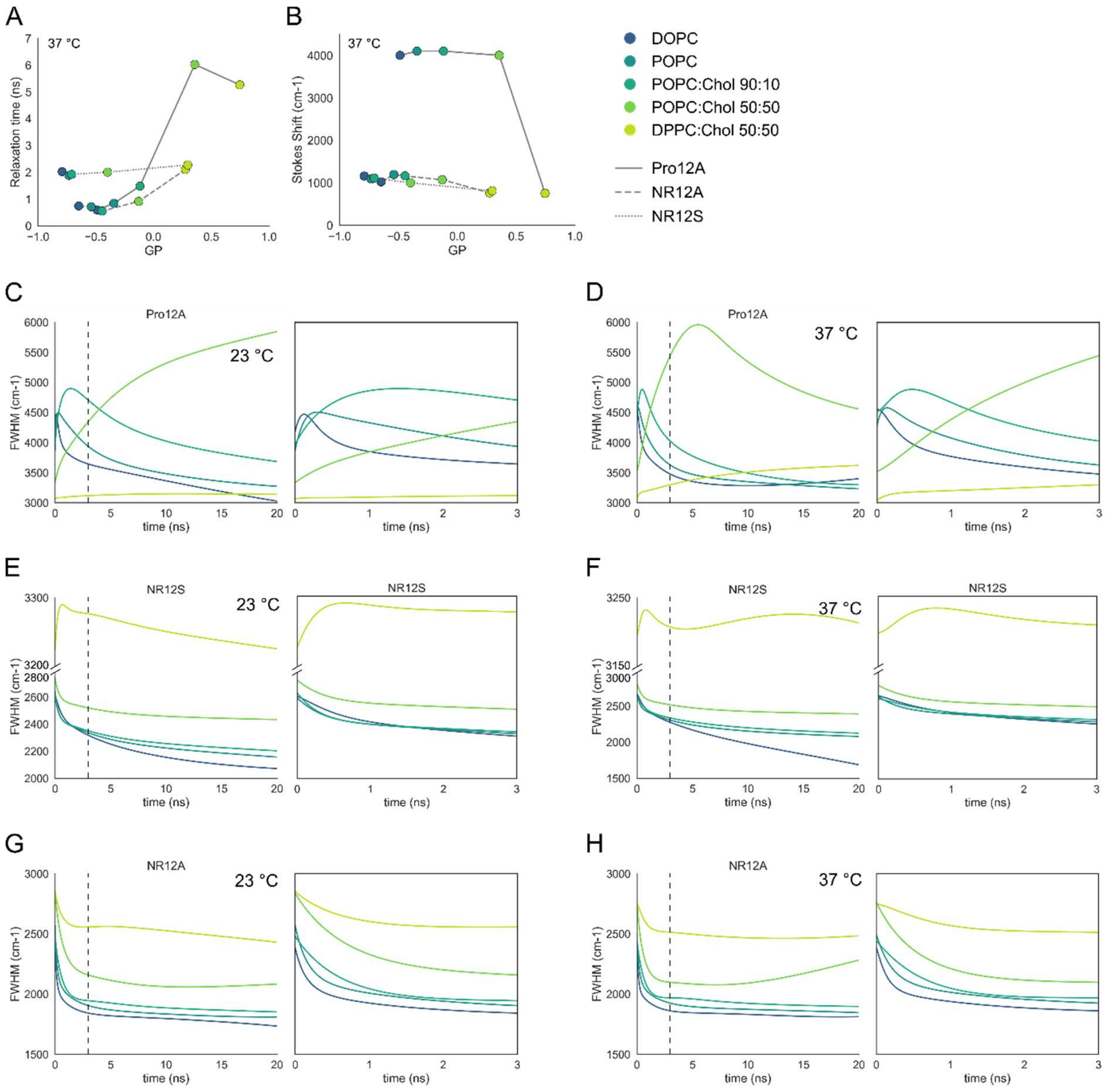


**Supplementary Figure 3: TDFS analysis of Pro12A, NR12S and NR12A reveals variations in relaxation behaviour in different lipid environments.**

TDFS analysis of the probes was performed in LUVs of varying lipid composition at 23 °C and 37 °C. Summary of the TDFS analysis of all probes can be found in Supplementary Tables 4-6. A| Relationship of Relaxation time and GP-value for all probes at 37 °C. B| Relationship of Stokes shift and GP-value for all probes at 37 °C. The line style indicates the probe. Colours indicate the different lipid compositions. C| FWHM of the TRES of Pro12A in different lipid compositions at 23 °C. Inset on the right shows FWHM from 0-3 ns. D| FWHM of the TRES of Pro12A in different lipid compositions at 37 °C. Inset on the right shows FWHM from 0-3 ns. E| FWHM of the TRES of NR12S in different lipid compositions at 23 °C. Inset on the right shows FWHM from 0-3 ns. F| FWHM of the TRES of NR12S in different lipid compositions at 37 °C. Inset on the right shows FWHM from 0-3 ns. G| FWHM of the TRES of NR12A in different lipid compositions at 23 °C. Inset on the right shows FWHM from 0-3 ns.

H| FWHM of the TRES of NR12A in different lipid compositions at 37 °C. Inset on the right shows FWHM from 0-3 ns. Colours indicate the different lipid compositions.
